# Adipose tissue in health and disease through the lens of its building blocks

**DOI:** 10.1101/316083

**Authors:** Michael Lenz, Ilja C.W. Arts, Ralf L.M. Peeters, Theo M. de Kok, Gökhan Ertaylan

## Abstract

**Background:** Highly specialized cells work in synergy forming tissues to perform functions required for the survival of organisms. Understanding this tissue-specific cellular heterogeneity and homeostasis is essential to comprehend the development of diseases within the tissue and also for developing regenerative therapies. Cellular subpopulations in the adipose tissue have been related to disease development, but efforts towards characterizing the adipose tissue cell type composition are limited due to lack of robust cell surface markers, limited access to tissue samples, and the labor-intensive process required to identify them.

**Results:** We propose a framework, identifying cellular heterogeneity while providing state-of-the-art cellular markers for each cell type present in tissues using transcriptomics level analysis. We validate our approach with an independent dataset and present the most comprehensive study of adipose tissue cell type composition to date, determining the relative amounts of 21 different cell types in 779 adipose tissue samples detailing differences across four adipose tissue depots, between genders, across ranges of BMI and in different stages of type-2 diabetes. We also highlight the heterogeneity in reported marker-based studies of adipose tissue cell type composition and provide novel cellular markers to distinguish different cell types within the adipose tissue.

**Conclusions:** Our study provides a systematic framework for studying cell type composition in a given tissue and valuable insights into adipose tissue cell type heterogeneity in health and disease.

## Background

The cell is the basic structural, functional, and biological unit of all living organisms. In multicellular organisms, a wide variety of highly specialized cells that work in synergy form tissues to perform functions required for the survival of the organism. The homeostasis in the system is maintained at the cellular level as well, such that defective, old, damaged or infected cells, or cells that are harmful to their environment either go through programmed cell death (apoptosis) or are actively detected and killed by other feedback mechanisms such as the immune system. Hence, many molecular sensors and inter-cellular mechanisms are evolved to ensure that the cellular heterogeneity is maintained at the tissue level.

Adipose tissue (AT) is no exception in this regard. It is composed of adipocytes, immune system cells, endothelial cells (blood and lymph vessels) and stem cells. Collectively, these cell types facilitate the functions associated with the tissue as an endocrine organ, energy depot, and major player in energy metabolism. To a large extent, it consists of adipocytes, which are commonly referred to as the fat depots in the body. Furthermore, adipose tissue has the unique ability to expand and shrink in significant proportions within the same individual over time. It can account for as little as 3% of total body weight in elite athletes or as much as 70% in morbidly obese individuals [1].

The conventional understanding of the adipose tissue portrays a fairly homogeneous tissue, responding to higher energy intake by expanding and to lower energy intake by shrinking. In contrast to this conventional belief, research in the last decade has focused on the different cellular subpopulations in the adipose tissue and their relation to (metabolic) health and disease [2,3]. This research provided deeper insights into disease development and progression in complex diseases (such as heart disease or diabetes) as well as inter-individual differences in disease etiology, paving the way for improved subtyping of patients for targeted therapies.

However, so far the efforts towards characterizing the adipose tissue cell type composition are limited, partially due to lack of robust cell surface markers identifying subpopulations of cells, but also due to limited access to tissue samples and the labor-intensive process required to identify them, such as immunohistochemistry or flow cytometry. Furthermore, the markers used to define a subpopulation of cells within the adipose tissue can differ greatly across studies, impeding reproducibility and leading to discrepancies across studies. Markers are usually defined for general purpose, and not designed to be specific for a tissue, bringing in to question their specificity and sensitivity.

In this paper, we propose a novel way of determining the adipose tissue cell type composition from whole tissue gene expression profiles. Our proposed TissueDecoder framework builds upon a recently published gene expression deconvolution algorithm [4], and facilitates reuse of published gene expression data for determining adipose tissue cell type composition across various depots and phenotypic traits (Figure 1). In this way, we are able to determine the relative fraction of 21 different cell types in 779 adipose tissue samples, presenting the most comprehensive study of adipose tissue cell type composition to date.

**Figure 1.**
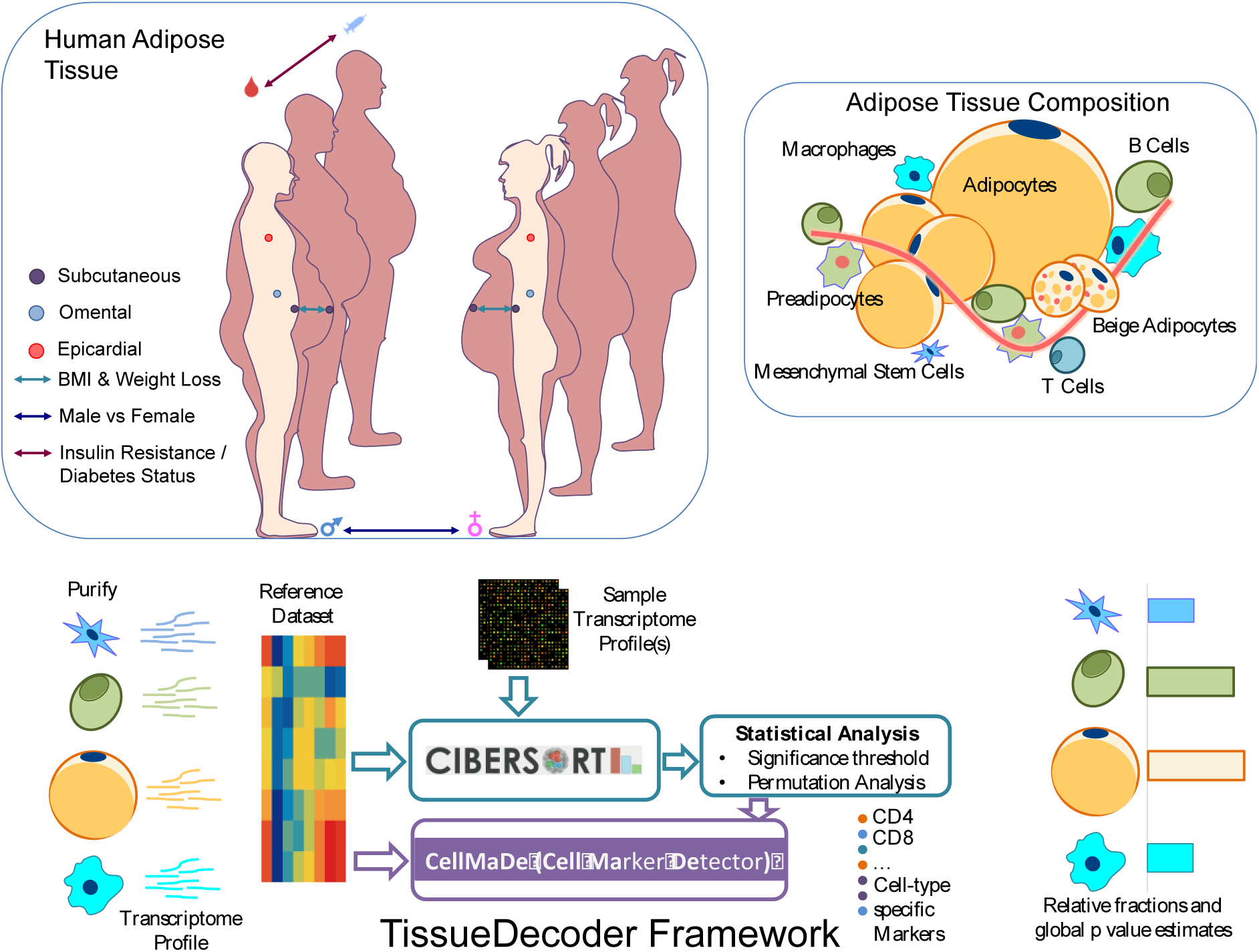
Visualization of the TissueDecoder framework for the analysis of adipose tissue cell type composition across a range of adipose depots and phenotypic traits. TissueDecoder consists of the proposed cell marker detector (CellMaDe) and the deconvolution of the cell types in adipose tissue using our AT21 signature matrix with CIBERSORT [4].

We determine the cell type composition of four different adipose tissue depots, including rare samples from epicardial (EAT) and pericardial adipose tissue (PAT), contextualize our results via a quantitative literature review and validate our approach using an independent dataset. Furthermore, we exemplify the usability of our approach by analyzing differences in adipose tissue cell type composition between genders, across ranges of BMI, and in different states of type 2 diabetes development.

Finally, we analyze the specificity of conventional cell type markers based on their gene expression level and introduce a novel algorithm (CellMaDe) for identification of cellular markers that are able to distinguish different cell types in a given tissue. Our study highlights the heterogeneity in reported studies of adipose tissue cell type composition and contributes to a better standardization by making our curated adipose tissue-specific signature matrix (AT21) available for future studies (Supplementary Data S1)

## Results

### Determining adipose tissue cell type composition from gene expression profiles

Our approach for determining the adipose tissue cell type composition from whole-tissue gene expression data employs the CIBERSORT deconvolution algorithm [4]. CIBERSORT uses a reference dataset of gene expression profiles from isolated cell types to generate a signature matrix, which is subsequently used as independent variable in nu-support vector regression (ν-SVR) to determine the cell type composition of whole-tissue samples. In this study, we extend the CIBERSORT signature matrix library with a novel signature matrix, termed AT21, since CIBERSORT is limited to blood due to its default signature. This broadens CIBERSORT’s applicability to deconvolution of the adipose tissue.

For our AT21 signature matrix, 21 cell types (Figure 2 A) from four cellular archetypes, namely immune cells (12 cell types), stem cells (adipose and (bone marrow-derived) mesenchymal), adipocytes (from subcutaneous adipose tissue (SAT) and PAT), and other cell types (endothelial cells, fibroblasts, smooth muscle cells, chondrocytes, and osteoblasts) were selected. We downloaded a total of 204 Affymetrix microarray datasets (Supplementary Data S2), containing samples from all 21 isolated cell types, from the gene expression omnibus (GEO) database [5], jointly preprocessed them, and generated the AT21 signature matrix using CIBERSORT [4] (see Methods).

**Figure 2.**
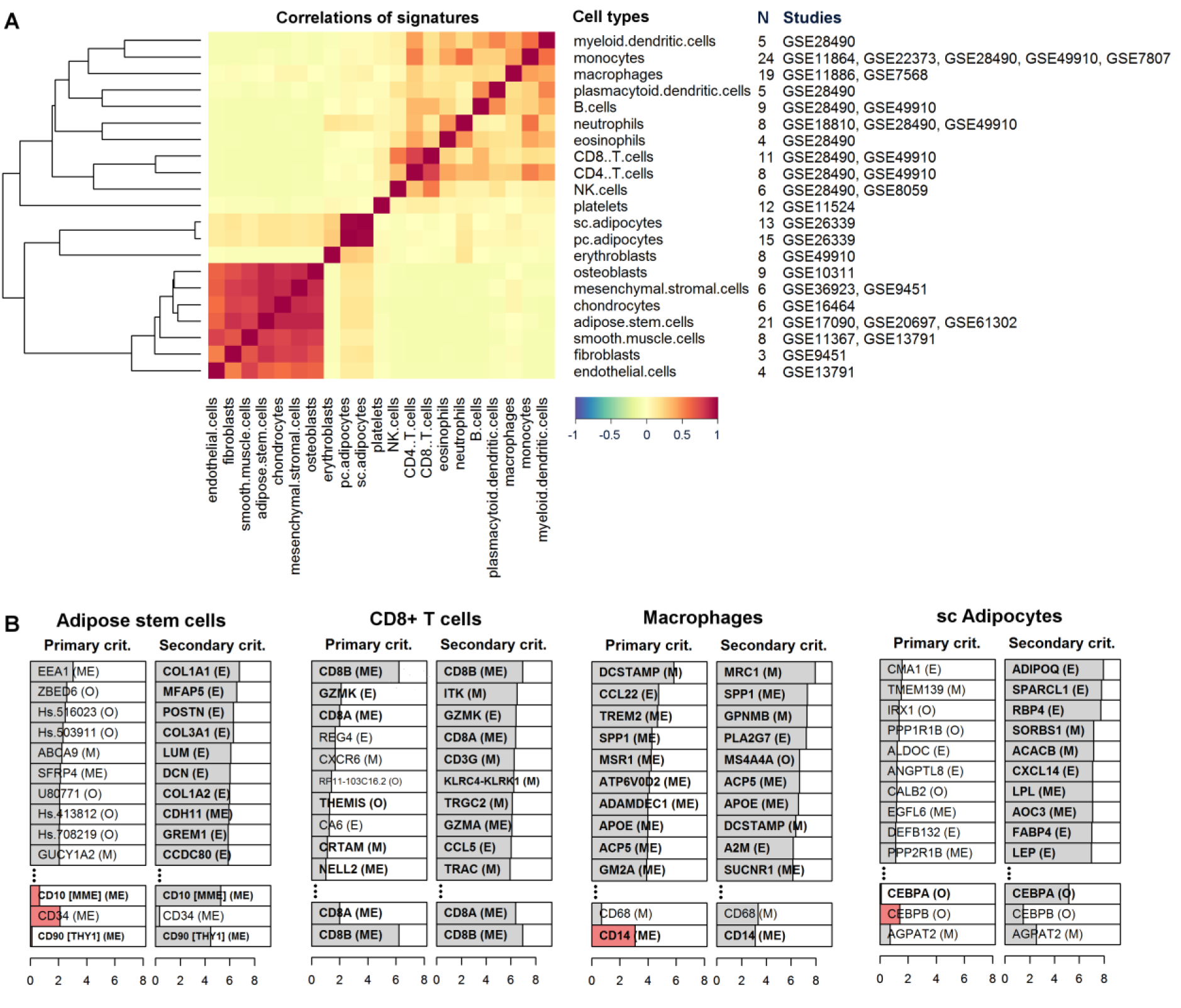
A) Heatmap showing the correlations of signatures for the reference dataset compiled from various studies reporting isolated cell type transcriptome profiles. The dendrogram (on the left) shows the clustering within the reference dataset. The dendrogram is constructed using hierarchical clustering with 1-correlation as the distance and average linkage as the linkage criteria. The number of samples contained for each cell type (N), and the GEO accession numbers per study is shown on the right. Three groups revealed higher correlation and clustered together in the reference dataset in accordance with our expectations: Highest correlation is observed between subcutaneous and pericardial adipose tissue followed by the cell types of the mesenchymal stem cell origin (osteoblasts, mesenchymal stromal cells, chondrocytes and ASCs) and finally the T cell group (CD4^+^ T and CD8^+^ T cell subsets) follow. B) Bar plots showing top ten cell type markers identified by the primary criterion and the secondary criterion of CellMaDe. Below each column conventional markers are demonstrated with their associated score. **Bold** font indicates the presence of the gene in the AT21 signature matrix and red colored bars indicate the value of the (primary/secondary) criterion was negative. Letters in brackets (M: membrane, E: extracellular, ME: both membrane and extracellular, or O: other) specify the gene ontology cellular location of the corresponding protein.

In Figure 2A we show the correlations of signatures between the different cell types, revealing high correlations between related cell types, such as subcutaneous and pericardial adipocytes, or mesenchymal stromal cells, chondrocytes, osteoblasts, and adipose stem cells (ASCs). In this analysis, we aim to evaluate the power of our approach to distinguish between similar cell types by following two strategies. First, we apply the deconvolution approach to the reference dataset itself, resulting in a clear distinction between cell types (Figure 3C). Second, we perform deconvolution analysis with a set of 779 adipose tissue samples (Supplementary Data S2) from four different adipose tissue depots (SAT, omental adipose tissue (OAT), PAT, and EAT) and check how well the results agree with the body of literature. This second analysis shows that the adipose tissue samples are predicted to have on average 14.5% ASCs, while the sum of the three related cell types (mesenchymal stromal cells, osteoblasts, and chondrocytes) is below 1% on average, revealing our ability to correctly distinguish between these cell types (Supplementary Figure S1). Furthermore, 95% of the SAT samples have a pericardial adipocyte score of 0% and an average subcutaneous adipocyte score of 74%, whereas OAT, EAT, and PAT samples are mostly predicted to have more pericardial adipocytes than subcutaneous adipocytes, although the distinction is less clear. This reveals the clear difference between subcutaneous adipocytes and adipocytes from other adipose depots in proximity to internal organs, which can be detected by the proposed methodology.

**Figure 3.**
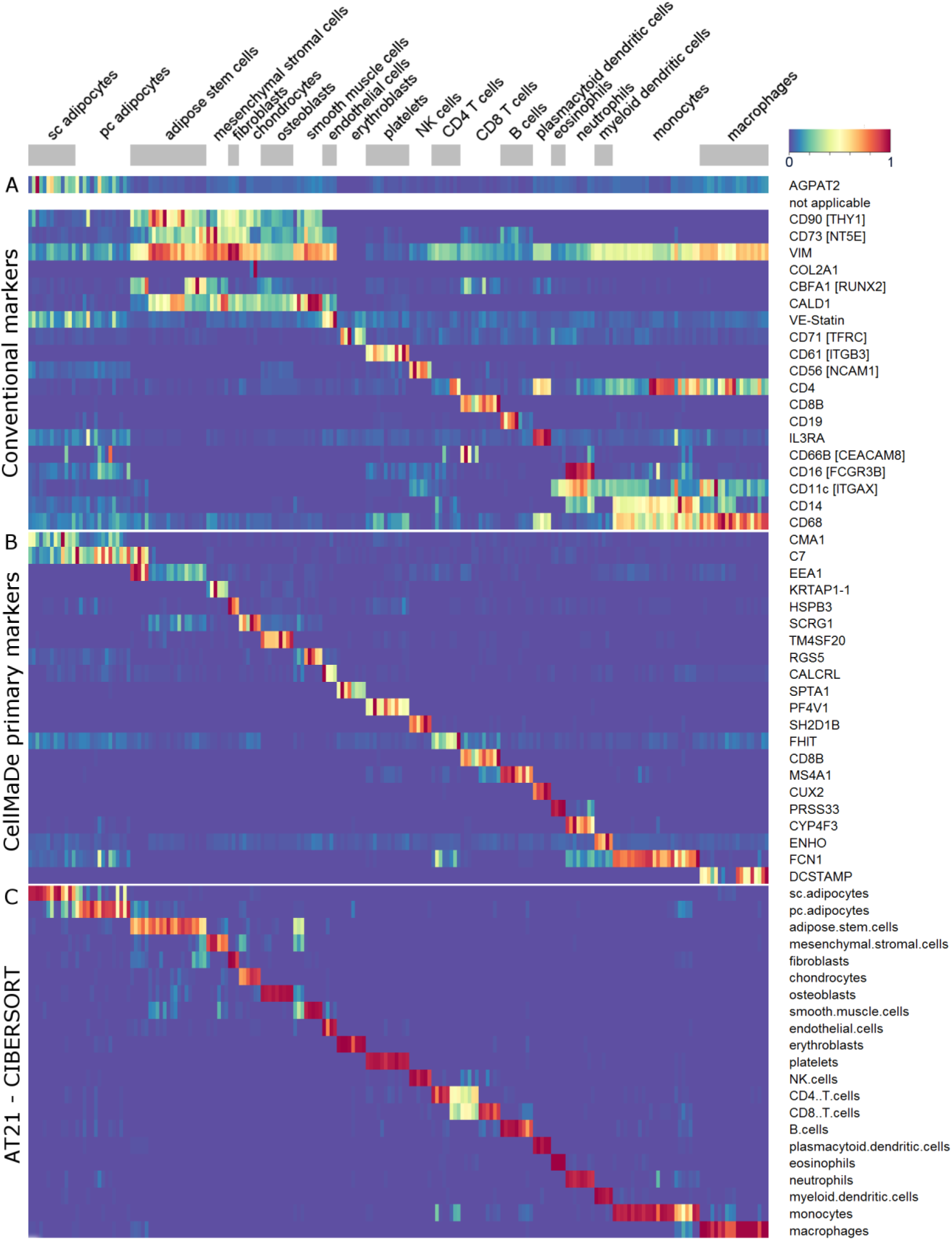
Heatmaps of AT21 signature matrix showing A) the normalized expression of selected conventional markers from literature, B) the normalized expression of CellMaDe predicted primary markers and C) cell type composition prediction from CIBERSORT with AT21 signature matrix. The common x-axis above the figure denotes the samples used for annotating cell types from the AT21 signature matrix. CellMaDe and CIBERSORT are both trained with AT21 signature matrix, therefore B) and C) are optimal results for both techniques whereas A) represents the separation power of conventional markers on the AT21 signature matrix.

### CellMaDe primary and secondary markers

Next, we further characterize the signature matrix by investigating which genes are incorporated in it, and by comparing their cell type specificity to that of conventionally used cell type markers. We have developed the *Cell Marker Detector* (CellMaDe) that uses two criteria to pinpoint i) highly specific markers that are only expressed in the target cell type and not in any other cell type of the tissue, referred to as *primary markers* (equation 1 in methods section), and ii) markers expressed in the target cell type that might also be expressed in some other cell types, referred to as *secondary markers* (equation 2 in methods section).

This has resulted in a ranking of all the probes present on the array to be annotated with their primary and secondary criterion scores (Supplementary Figure S2, Supplementary Data S3). In Figure 2B, we show the top 10 genes having the highest primary and secondary criterion score from four cell types - namely ASCs, CD8^+^ T cells, macrophages and subcutaneous adipocytes – and indicate their cellular location according to gene ontology terms. All genes that are present in the AT21 signature matrix are marked in bold, showing that CIBERSORT algorithm mainly selects a combination of secondary markers (CIBERSORT’s gene selection criterion is comparable to our secondary marker criterion), while primary markers and conventional markers are not always included in the signature matrix.

This is in contrast to classical marker-based approaches such as immunohistochemistry, which rely heavily on the cell type specificity of a single (primary) marker. Nevertheless, conventional markers such as CD68 for macrophages are also expressed in other cell types, such as monocytes, plasmacytoid dendritic cells [6], and to a lesser degree also in fibroblasts and endothelial cells [7,8], which is also indicated by its relatively low primary criterion score (Figure 2B). We, therefore, propose the primary markers identified by CellMaDe to be further validated experimentally and used as alternative markers for the cell types within the adipose tissue.

For CD8^+^ T cells, the conventionally used marker (Cluster of differentiation 8B; CD8B) is identified as the strongest primary marker, which is not very surprising. However, Cluster of differentiation 4 (CD4) is not very specific for CD4^+^ T cells, as shown by its prevalent expression in other hematopoietic cell types such as monocytes or macrophages (Figure 3A, Supplementary Figure S2). The Dendrocyte Expressed Seven Transmembrane Protein (DCSTAMP) is identified as a very specific primary marker for macrophages according to our analysis. In Figure 3B we observe that the high expression of DCSTAMP is limited to a subset of the macrophage samples, which might highlight its specificity to a subset of macrophages, e.g. macrophage giant like cells, as suggested by earlier studies [9]. For ASCs and subcutaneous adipocytes, the identified primary markers (EEA1 and CMA1, respectively) are not very striking. For these cell types, we suggest using a combination of secondary markers as implemented in flow cytometry gating strategies as well as in CIBERSORT algorithm.

For further evaluation of the identified top primary markers, we compared their expression in a more extensive independent dataset comprised of 394 anatomically annotated tissue expression profiles (Supplementary Figure S3, Methods). Six of the 21 primary markers are fully validated, defined as showing the highest expression in the proposed cell type, namely CALCRL for endothelial cells, SPTA1 for erythroblasts, CD8B for CD8^+^ T cells, MS4A1 for B cells, PRSS33 for eosinophils, and DCSTAMP for macrophages. Twelve markers are partially validated, identified as being among the top five of the 394 anatomically annotated tissue expression profiles or being expressed at a higher level only in cell types that are not related to adipose tissue, such as myocytes or neurons. Only three markers were not validated, namely EEA1 for ASCs, RGS5 for smooth muscle cells, and PF4V1 for platelets, the latter due to the absence of platelets in the validation dataset for PF4V1, preventing its proper evaluation.

### Independent (ex-vivo) validation

We have performed an independent validation of our approach using a “validation dataset” (see methods) including adipocytes, progenitor cells (CD45^-^CD34^+^CD31^-^) and monocytes/macrophages (CD45^+^CD14^+^) directly isolated from adipose tissue across 19 individuals (10 control and 9 obese) [10] and B cells, CD4^+^ T cells, CD8^+^ T cells, natural killer cells, and monocytes directly isolated from blood [11]. We use this validation dataset to evaluate the gene expression deconvolution approach on isolated cell types and to test the specificity of the proposed primary markers identified by CellMaDe.

The results of this analysis are presented in Supplementary Figure S4 and Supplementary Data S2, showing that the primary markers are expressed and able to distinguish between cell types in the validation dataset, with the exception of CMA1 for subcutaneous adipocytes and EEA1 for ASCs (Supplementary Figure S4 B). The CIBERSORT-based deconvolution with the AT21 signature matrix correctly provides highest percentages for the respective isolated cell type for all tested cell types in the validation dataset (Supplementary Figure S4 C).

The mean estimated percentages of the isolated cell type are 69.2% for subcutaneous adipocytes, 57.1% for ASCs, 89.9% for B cells, 84.8% for CD4^+^ T cells, 71.0% for CD8^+^ T cells, 62.0% for NK cells, and 90.5% for monocytes. The CD14^+^ fraction of the adipose tissue (monocytes/macrophages) is predicted to consist mainly of macrophages (25.4%), myeloid dendritic cells (24.4%) and monocytes (18.1%). The deviation from the optimal results based on the reference data itself (Figure 3 C) can be explained by cross-platform differences between the validation and reference datasets. (the validation dataset and reference dataset have been hybridized to two different Affymetrix microarrays).

### Contextualization of findings – a literature review

As the next step of evaluation, we have compiled quantitative literature reports of human adipose tissue cell type composition to compare the estimates of our deconvolution approach for 616 SAT samples and 51 OAT samples to the reported percentages in the literature.

The first study that aimed to systematically categorize different cell types in adipose tissue from human samples used flow cytometry on liposuction aspirates to quantify different types of cells present in the adipose tissue [12]. Our literature search resulted in 25 studies reporting cell counts for 10 different cell types, including macrophages (18 studies), CD4^+^ and CD8^+^ T cells (5 studies), B cells (4 studies), Natural killer cells (2 studies), myeloid dendritic cells (1 study), adipocytes (1 study), ASCs (9 studies), endothelial cells (5 studies), and fibroblasts (1 study). In order to make the reported results comparable, we converted the units – cells/100 adipocytes, % of stromal vascular fraction (SVF), cells/total number of nuclei, or cells/g AT – into percent of total cells in the adipose tissue (methods).

The results of the literature review are presented in Figure 4 and Supplementary Data S4. Reported cell counts for macrophages vary greatly, ranging from average counts of less than 1% of total cells in some studies up to an average of 27% of total cells in another study. These rather large differences between the studies can arise due to several factors, including (i) actual biological differences between the analyzed samples, (ii) local enrichment of macrophages (e.g. in crown-like structures) that specifically influence results with low total cell counts like immunohistochemistry, (iii) technical differences between the utilized methodologies and markers, (iv) differences in sample handling and analysis protocols (e.g. fluorescence cutoffs, utilized antibodies), and (v) differences in reported units across different studies.

**Figure 4.**
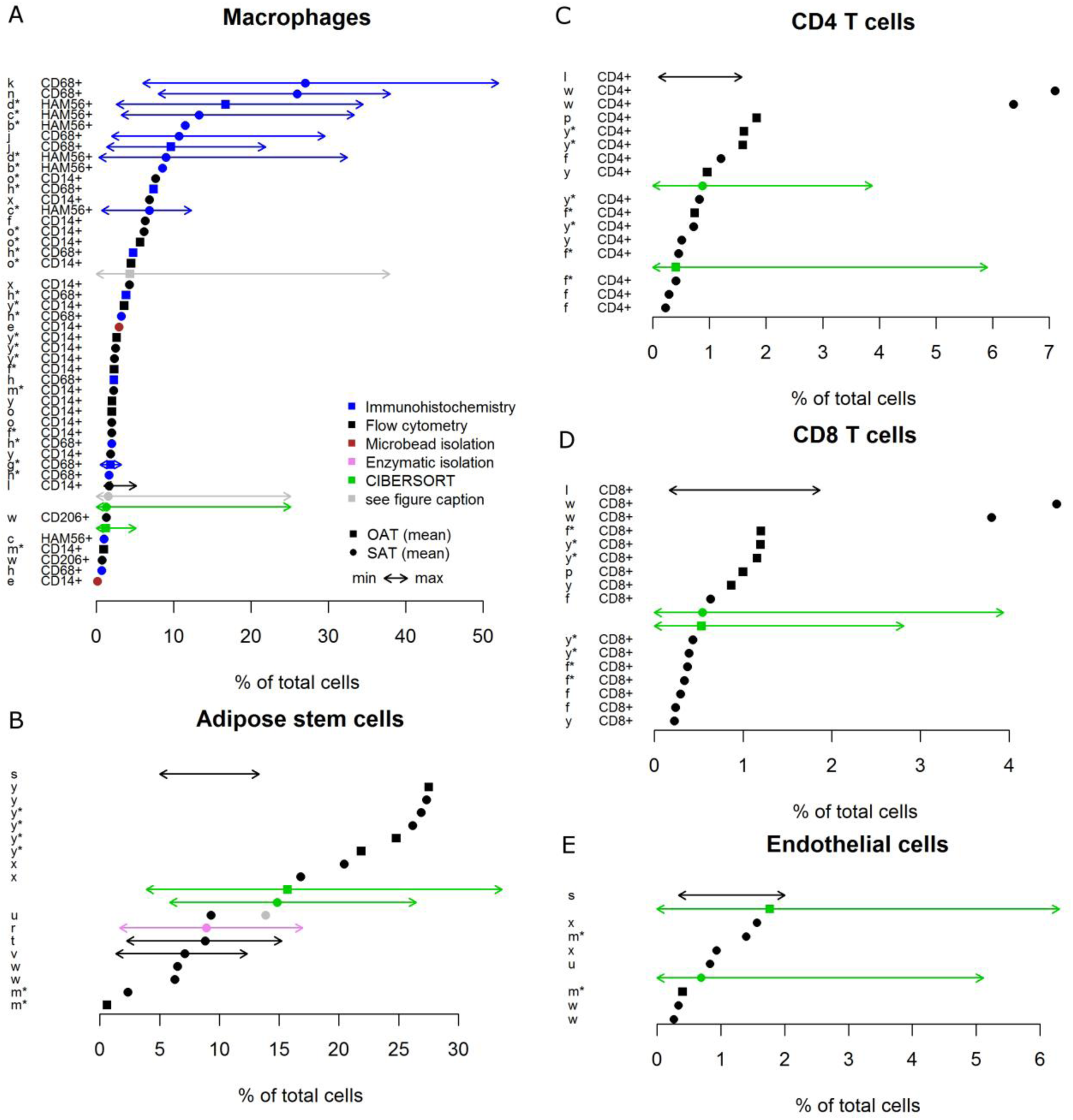
Literature review of reported adipose tissue cellular composition in comparison to our results using CIBERSORT (in green). Shown is the mean (dots), minimum, and maximum (arrows) percentage of total cells for five different cell types (macrophages, ASCs, CD4^+^ T cells, CD8^+^ T cells, and endothelial cells). The color specifies the method used for cell counting as indicated in the legend. The gray dots and arrows in (A) are our results where the macrophage score and monocyte score have been added together. It is shown as a comparison, since the macrophage markers CD68, HAM56, and CD14 also stain monocytes. The gray dot in (B) represents the sum of ‘supra adventitial-adipose stromal cells’ (black dot in the same row) and ‘endothelial progenitor cells’, which were distinguished in the respective study, but are likely both covered in the ‘adipose stem cell’ score from our AT21 signature matrix. On the left hand side of each plot the references to the studies from which the results were taken (see Supplementary Data S4) and the utilized markers are indicated. For ASCs and endothelial cells a combination of markers was used (see Supplementary Data S4) and the CD206 marker in (A) was used in combination with CD14. A star attached to the study reference letter indicates that the study participant had an average body mass index above 35. It is included in the figure since it has been reported that the macrophage frequency is increased in people with severe obesity.

It is important to note that specifically, some immunohistochemistry studies report very high macrophage numbers (Figure 4A). In addition, there may be some differences due to reported units across different studies, as the two studies with highest macrophage percentages (averages of 27% and 26% macrophages) are the only ones reporting in ‘macrophages per total number of nuclei’. We have shown previously that immunohistochemistry studies from tissue slices can be biased due to reliance on observations from cross-sections (thin tissue slices) [13]. Therefore, it can be argued that specifically for adipocytes the cross-section may cover parts of the lipid droplet, but not the nucleus, resulting in systematic differences between counting methodologies.

In order to evaluate the potential influence of biological differences between study participants, especially with respect to their obesity status, we marked all studies involving people with an average body mass index above 35 (severe obesity) with a star (Figure 4A). The relationship between obesity status and macrophage counts has been studied in several articles, reporting increased macrophage counts with increasing obesity in some, but not all studies [14-19]. Nevertheless, the obesity status cannot explain the observed diversity between reported macrophage percentages in our literature review (Figure 4A).

For comparison, we assessed inter-study differences in our analyses (only SAT), showing relatively stable results, which indicates a better standardization despite biological differences of study participants and potential differences in sample handling between different labs (Supplementary Figure S5).

In comparison to the literature reports, the estimated amount of macrophages from our analysis is rather at the lower end, with an average of 1.3% of total cells for the 616 SAT samples (median of 0.8%, IQR: 0.03%-1.8%) and 1.2% of total cells for the 51 OAT samples (median of 1.2%, IQR: 0.4%- 1.8%). Looking at the extreme values, our results confirm that there is a large range of macrophage frequencies with up to 25% macrophages in very rare cases.

In order to account for the potential influence of monocytes, which also express the markers utilized in the literature studies (CD14, CD68, and HAM56), we also report the combined fractions of macrophages and monocytes from our AT21-CIBERSORT approach. This amounts to an average of 1.5% macrophages/monocytes in SAT samples and 4.3% macrophages/monocytes in OAT samples, which brings our estimates in close proximity to the values reported in the literature.

The amount of endothelial cells (mean of 0.7% in SAT and 1.8% in OAT), CD4^+^ T cells (mean of 0.9% in SAT and 0.4% in OAT), CD8^+^ T cells (mean of 0.5% both in SAT and OAT) and ASCs (mean of 14.8% in SAT and 15.7% in OAT) estimated by our approach is well in line with the literature reports. Similar to macrophages, literature reports of ASC quantities show large variations (Figure 4E). We have identified three reasons explaining this variation. First, three of the literature studies (studies s, t, and v – see Supplementary Data S4) use adipose tissue from liposuction aspirates, which is contaminated with blood [12], resulting in lower relative fractions of ASCs in the SVF. Second, some studies distinguish between endothelial progenitors and supra-adventitial adipose stem cells (e.g. studies u and v), whereas others (including our study) don’t, possibly counting both subpopulations as ASCs. Notably, when adding up the two subpopulations in the study of Zimmerlin [20], the resulting average fraction of 13.9% (gray dot in Figure 4E) is close to our estimated results. Third, the total cell yield and stem cell percentage vary widely depending on the method used for isolation of SVF [21,22], which can potentially explain the very low numbers reported by Viardot and colleagues [23] (study m).

There are several cell types such as plasmacytoid dendritic cells, neutrophils, or eosinophils for which we could not find any results in the literature and thus we are reporting their frequency in the human adipose tissue for the first time. Some cell types such as platelets and erythroblasts are mainly included as a control in the AT21 signature matrix, allowing for identification of the presence of blood within the samples.

### Comparison of four adipose tissue depots

Next, we compare the cell type composition of four adipose tissue depots (SAT, OAT, PAT, and EAT) by reporting their average cell type composition (Figure 5, detailed in Supplemental Figure S1). This indicates that SAT has the highest percentage of adipocytes (74%) followed by OAT (66.4%), EAT (59.5%) and PAT (59.4%), while EAT and PAT have far more immune cells (20.8% and 20.9%, respectively) compared to OAT (9.8%) and SAT (7.4%). Furthermore, OAT is the richest source of stem cells (17.2% compared to 14.9% for SAT, 14.1% for EAT and 12.4% for PAT).

**Figure 5.**
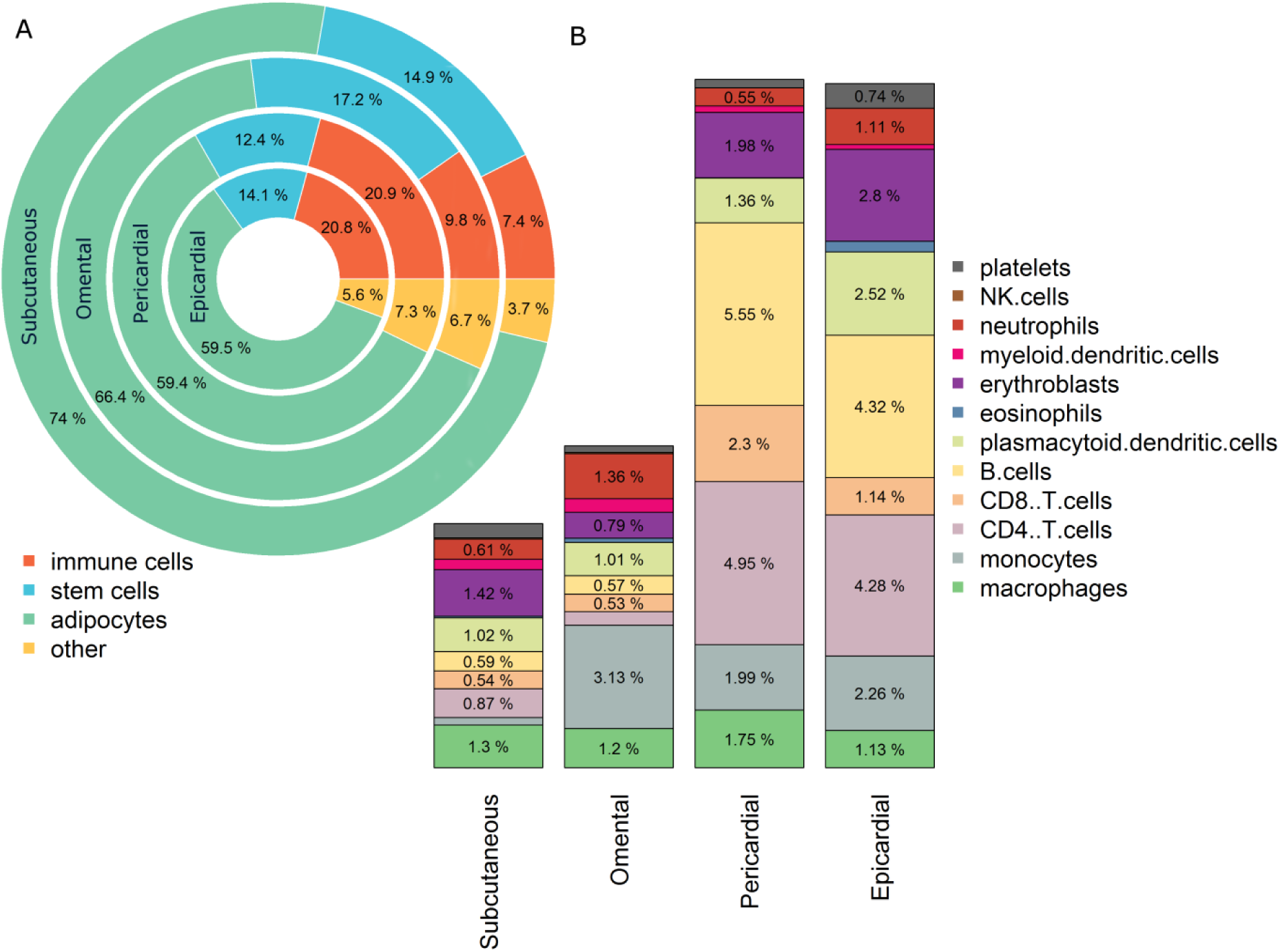
Estimated percentage of cell types per adipose depot. (A) Pie chart showing the overall cell type distribution of various adipose tissue depots (Subcutaneous – n=616, Omental – n=51, Pericardial – n=66, and Epicardial – n=46) in terms of four main archetypes of cells in adipose tissue (Immune cells, Stem cells, Adipocytes, and other). (B) Bar plots showing the detailed distribution of the immune cell compartment from each adipose tissue depot. All values are averages of the analyzed samples from the respective depot.

These results need to be interpreted with care due to differences in number and characteristics of people from which the samples were collected. Access to EAT and PAT is severely limited due to their physiological location and the invasive nature of the sampling procedure. Therefore PAT samples were taken from 66 patients (age: 66±8 years) with coronary artery disease (CAD) [24] and the EAT samples were taken from 11 neonates (6 to 24 days old), 28 infants (40 days to 1 year-old) and 7 children (2 to 7 years old) with congenital heart disease (CHD) [25].

Despite differences in the age of EAT and PAT donors, the composition of the immune cell archetype in EAT and PAT is remarkably similar to each other while being very different from SAT and OAT samples. This indicates the robustness of our results as well as the conserved nature of the cell type composition in these two adipose tissue depots surrounding the heart.

Furthermore, we report the fractions of classically designated “adaptive” immune cells, including B cells, CD8^+^ and CD4^+^ T cells, infiltrated into EAT and PAT. These adaptive immune cells are enriched in EAT and PAT, (higher in PAT than in EAT) compared to SAT and OAT depots, which could likely be due to the origin of the analyzed samples from patients with CAD or CHD, as reported earlier for CD8^+^ T cells [26]. Our finding is also supported by Mazurek and colleagues [27], who compared the expression of cytokines in both EAT and SAT in patients undergoing coronary artery bypass graft and found that EAT is a source of several inflammatory mediators.

A more detailed comparison of SAT and OAT was performed on the study of Hardy et al 2011 (dataset GSE20950), in which both depots were available from the same individuals [18]. This analysis revealed an increased neutrophil content in SAT (also after correcting for multiple testing) and increased mesenchymal stromal cell and smooth muscle cell content in OAT (Figure 6A).

**Figure 6.**
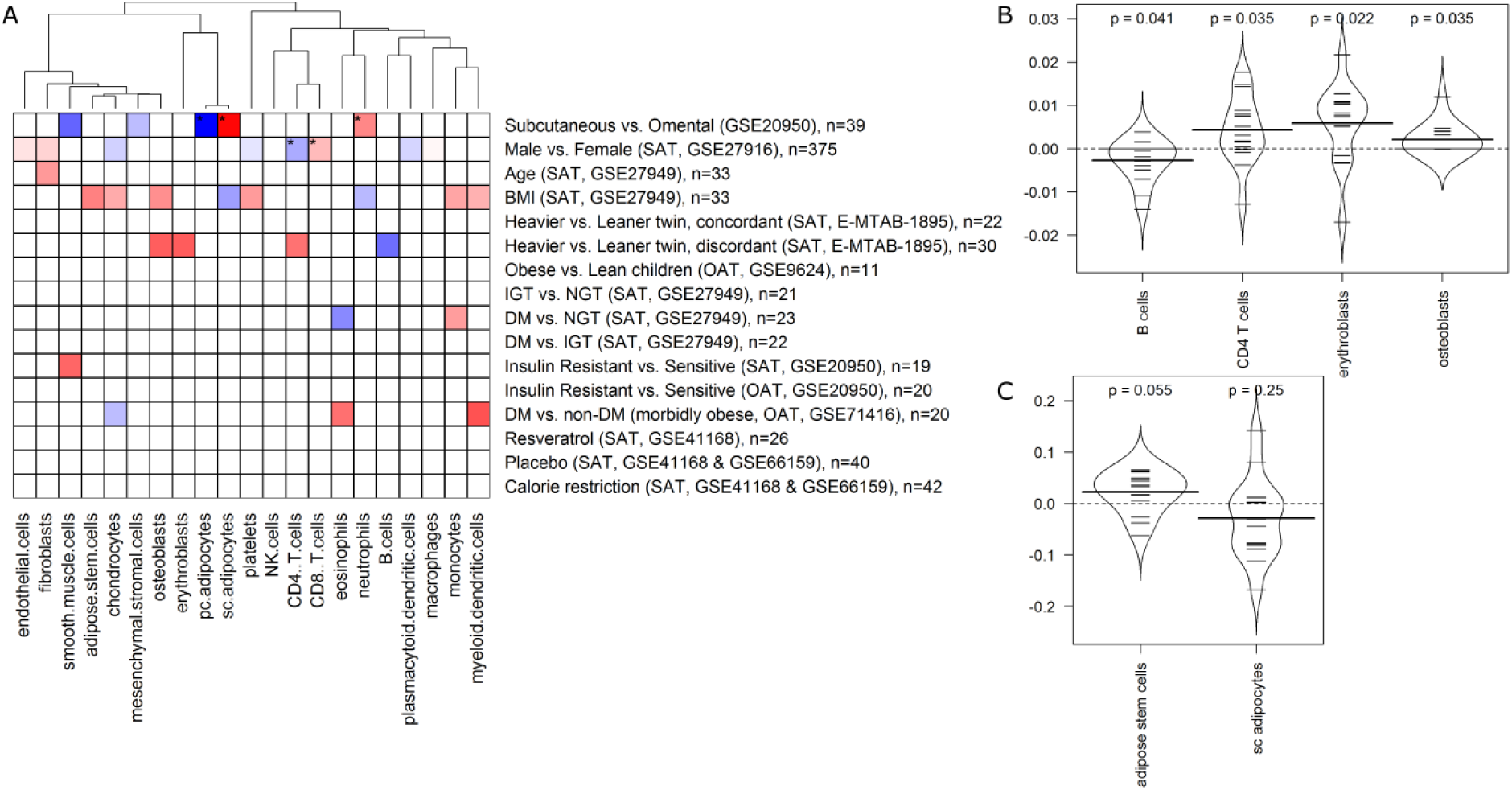
A) Heatmap showing the significant over (red) /under (blue) represented cell type across different categories based on the z-score (denotes the standard deviation each group is away from the other). Only significant results (uncorrected p<0.05) are colored. Stars label results that stay significant after correction for multiple testing. The bean plots showing B) all significant cell types and C) ASCs and subcutaneous adipocytes (not detected as significant), with the population distribution of the cell type percentages for the heavier vs leaner twin, discordant study. The y axis describes differences in estimated cell fractions between the heavier and leaner twin within a twin pair.

### Association of cell type composition with phenotypic traits

Finally, we compared the adipose tissue cell type composition between individuals with different phenotypic traits, such as different gender, age, body mass index (BMI), and different stages of type 2 diabetes development. Furthermore, we included intervention studies investigating the effect of caloric restriction or resveratrol ingestion on SAT cell type composition. The results of these analyses are shown in Figure 6 and in more detail in Supplementary Figure S6 and Supplementary Data S5.

We detected significant differences in SAT cell composition between males and females, indicating that there are higher amounts of plasmacytoid dendritic cells, CD4^+^ T cells, platelets, and chondrocytes in females, whereas males have more CD8^+^ T cells, fibroblasts, and endothelial cells. Notably, the differences in CD4^+^ and CD8^+^ T cells were significant also after correction for multiple testing. With respect to age, we only detected one significant result, indicating a higher amount of fibroblasts with increasing age.

We include four comparisons related to BMI, namely (i) a continuous relationship of BMI with cell fractions in SAT, (ii) a comparison of SAT cellular composition in concordant monozygotic twins (heavier vs. leaner twin, ∆BMI < 3 kg/m^2^), (iii) the same in discordant monozygotic twins (heavier vs. leaner twin, ∆BMI > 3 kg/m^2^), and (iv) a comparison of OAT cellular composition in obese vs. lean children. The first comparison indicates that BMI is positively correlated with the amount of monocytes, myeloid dendritic cells, platelets, osteoblasts, chondrocytes, and ASCs in SAT, whereas the correlation with subcutaneous adipocytes is negative. No significant differences are detected between concordant twins, whereas discordant twins differ significantly in the amount of CD4^+^ T cells, erythroblasts, osteoblasts (all higher in heavier twins), as well as B cells (higher in leaner twins). Notably, there is a tendency towards higher amounts of ASCs and lower amounts of adipocytes in heavier twins (Figure 6C), supporting the respective significant finding in the continuous association with BMI. In contrast, some other findings of the continuous association with BMI are not detected in the twin study, which can be attributed to potential confounding effects (age, gender, or other factors) in the continuous association.

The lack of any significant results in the comparison of lean vs. obese children is mostly due to limited power since the study only involves a total of 11 samples (5 obese and 6 lean children).

We made six comparisons related to diabetes, glucose tolerance, or insulin resistance. Three of them (impaired glucose tolerance vs. normal glucose tolerance in SAT; diabetes mellitus vs. impaired glucose tolerance in SAT; and insulin-resistant vs. sensitive in OAT) did not show any significant results. The comparison of SAT cell type composition in insulin-resistant vs. sensitive individuals revealed a significant increase in smooth muscle cells in insulin resistant individuals. The SAT of people with diabetes mellitus contains more monocytes and fewer eosinophils than that of normal glucose tolerant people, according to our analysis, whereas the OAT of morbidly obese people shows differences in myeloid dendritic cells, eosinophils (both higher in people with diabetes), and chondrocytes (higher in people without diabetes).

We did not find any evidence that caloric restriction or resveratrol supplementation significantly changes adipose tissue cell type composition.

Interestingly, one included study also reports macrophage percentages as determined by immunohistochemistry [18] (GSE20950). They show macrophage frequencies for 11 OAT samples (out of 20 study participants) from insulin sensitive and insulin resistant people, with an average macrophage content of 1.8% (after unit conversion to % of total cells), which nicely matches our estimate of 1.495% as average of all 20 participants (selection criteria of the 11 samples for immunohistochemistry were not included in the study description). They furthermore report a significant association between HOMA-IR and macrophage content in these 11 people, which we partially confirm with borderline significance (p-value of 0.054) for a comparison of insulin-resistant versus insulin sensitive people in all 20 study participants.

## Discussion

Tissues consist of numerous different cell types and this heterogeneity plays a prominent role in tissue homeostasis in health and disease. In this work, we propose the cell type composition of a tissue, specifically adipose tissue, as a way to investigate the differences across individuals as well as biological conditions.

Here we describe our novel findings on the cell type composition of the human adipose tissue. We compare our findings to the existing body of knowledge attained via a range of methods (immunohistochemistry, flow cytometry, microbead isolation and enzymatic isolation) by including a review of the studies addressing quantification of cell types (macrophages, CD4^+^ T cells, CD8^+^ T cells, endothelial cells and ASCs) within the subcutaneous and omental adipose tissue. The meta-analysis of these studies indicates that, with the exception of macrophages and adipose tissue stem cells (ASCs), there is clear consensus on the extent of the presence of the above-mentioned cell types within the adipose tissue, which clearly overlaps with the results of our analysis. It is postulated that macrophages constitute a specific population of cells that can differ greatly due to inter-individual differences. The extent of tissue specificity for macrophages is reviewed elsewhere [28].

ASCs are considered to be highly suitable for a range of applications in regenerative therapies [29-31]. Despite all the prospects, our understanding of the ASC within the adipose tissue is currently quite limited. The estimates of ASCs within the adipose tissue are varying from 1-27 % of the total cells (Figure 4B) depending on the exact markers used to define the ASC phenotype as well as the type of method used for isolating ASCs. In this study we robustly define the ASC signature based on twenty one samples from three independent datasets and report that the ASC composition within the adipose tissue can greatly vary from 3% to over 30% across individuals with the mean of 14.8% in SAT (IQR: 12–17.2%) and 15.7% in OAT (IQR: 11.5–18.6%). This relatively large variation is potentially due to the dynamic nature of the stem cell compartment as well as the broad spectrum of individuals selected for this study. These results highlight adipose tissue as the richest source of multipotent stem cells, outnumbering possibly any other source in the body.

Moreover, we report for the first time gender-specific differences in the immune system compartment within the subcutaneous adipose tissue, with clear enrichment of CD4^+^ T cells in females and enrichment of CD8^+^ T cells in males. This is in line with the established notion of higher numbers of CD4^+^ T cells and CD4^+^/CD8^+^ T cell ratios in females versus males observed from the peripheral blood mononuclear cell (PBMC) compartment. This gender difference is reported to remain constant from birth to old age and has potential consequences for a range of conditions including susceptibility to infectious diseases as well as risk for autoimmunity and non-reproductive cancers [32]. The role of gender-specific differences in adipose tissue CD4^+^/CD8^+^ T cell ratios is yet to be further investigated.

We also compared the cell type composition of different adipose tissue depots, revealing high stem cell percentages (12.4%-17.2%) in all depots, identifying SAT as richest in adipocytes (74%) and EAT and PAT as highly enriched in immune cells, 20.8%, and 20.9% respectively. Our results implicate robust and systematic differences in cell type composition across adipose tissue depots adding to previously identified differences in metabolism and adipocyte size between depots [33].

The results highlighted above demonstrate i) cell type composition differences across adipose tissue depots, revealing adipose tissue heterogeneity and assisting future studies investigating the role of a specific adipose tissue depot with a disease phenotype; ii) gender-specific differences in CD4^+^/CD8^+^ T cell ratios within the subcutaneous adipose tissue providing a completely new angle and a potential confounding factor while studying metabolic syndrome; iii) adipose tissue as the richest source of multipotent stem cells underlining the significance of adipose tissue in the stem cell field, paving the way for future regenerative therapies. Taken together, they demonstrate the power of the TissueDecoder framework in enumeration of cell types within adipose tissue and give an overview of its potential in relating to other fields.

Our approach has various advantages including its ability to analyze many cell types in parallel, its relatively high level of standardization, and its compatibility with retrospective datasets, allowing for meta-analyses of public data from various laboratories. Furthermore, it has the potential of being platform (e.g. Affymetrix or Illumina microarrays) and technique (microarray or RNA-Seq) independent, although most accurate results are currently produced by using the same platform for the reference dataset and analysis dataset, as employed in our study.

Naturally, the methodology proposed also comes with limitations that can be addressed in future versions or by other studies. One potential extension would be the inclusion of brown/beige adipocytes in the reference dataset in conjunction with an inclusion of adipose tissue depots harboring brown adipose tissue such as supraclavicular, paravertebral, mediastinal, para-aortic and suprarenal depots.

Furthermore, many cell types in the AT21 dataset were not extracted from adipose tissue. Generating a reference dataset in which all cell types are isolated from the adipose tissue could potentially increase the accuracy of the method since for instance macrophages are known to have distinct expression depending on their tissue of origin [28]. Related to this, the AT21 reference dataset has been generated by reference cell types that were isolated based on a few cell surface markers, which in theory could potentially bias the identification. We propose to use single-cell transcriptomics analysis in future studies to identify the cell types for generation of the reference sets [34].

Finally, in order to study the relation of phenotypic traits with adipose tissue cell type composition in more detail, it is necessary to include studies with larger sample sizes, possibly a few hundred samples, which in our case was only available for the comparison of male vs. female.

## Conclusion

In conclusion, we report a novel methodology to quantify the tissue cell type composition in adipose tissue that can be used both in future studies as well as in existing datasets with our established signature matrix. In generalized term, our framework allows tissue cell type composition to emerge as a potential marker measuring homeostasis that can be utilized in prospective studies in regenerative medicine. The overview of studies presented in comparison with our findings, reflect the state of the adipose tissue field from the lens of tissue cell type composition. Finally, reported novel cell type-specific markers have great potential in identifying difficult cell types (such as adipose tissue macrophages and eosinophils) in future studies working with adipose tissue samples.

## Methods

### TissueDecoder Framework

#### Deconvolution of adipose tissue cell types

In the TissueDecoder framework, CIBERSORT [4] is being used to determine the cellular composition of adipose tissue samples from their whole tissue gene expression profiles. CIBERSORT uses a signature matrix of cell type-specific gene expression in combination with a deconvolution approach based on nu-support vector regression. The focus of the original publication is on immune cell types from the blood. Therefore, CIBERSORT’s default signature matrix does not include adipocytes, endothelial cells, or ASCs, limiting its applicability to be used in other tissue types. In order to apply CIBERSORT for the deconvolution of adipose tissue, we have curated a novel signature matrix (AT21) that includes highly relevant cell types for adipose tissue. We have generated the AT21 signature matrix based on publicly available data from the Affymetrix Human U133 Plus 2.0 microarray (GEO identifier ‘GPL570’). We restricted our search to this microarray platform to avoid compatibility problems with cross-platform differences.

##### Reference dataset and AT21 signature matrix

For the generation of AT21, we collected single cell type gene expression data from 21 different cell types (204 samples in total) from publicly available datasets in the Gene Expression Omnibus (GEO) database [35] (Figure 2A). Raw data (CEL files) were downloaded and preprocessed with Affymetrix Power Tools (https://www.thermofisher.com/nl/en/home/life-science/microarray-analysis/microarray-analysis-partners-programs/affymetrix-developers-network/affymetrix-power-tools.html#) using the robust multi-array average (RMA) normalization method.

The normalized reference dataset was then used to generate the AT21 signature matrix using CIBERSORT [4] (https://cibersort.stanford.edu). For each cell type, CIBERSORT first filters probes based on their differential expression between the selected cell type and all other samples (q value<0.3 (false discovery rate), two-sided unequal variance t-test). Subsequently, probes are ranked according to their fold change between the respective cell type and all other samples and the top G probes are included in the signature matrix. Here, G (between 50 and 150) is selected to minimize the condition number of the signature matrix [4]. This resulted in a total of 1872 probes in AT21 (Supplementary Data S1).

##### Adipose tissue samples and deconvolution

We utilized the AT21 signature matrix to determine the cell type composition of 779 adipose tissue samples hybridized to the same microarray platform as the reference dataset (Affymetrix Human U133 Plus 2.0). The adipose tissue samples came from four different adipose depots (subcutaneous, n=616; omental, n=51; pericardial, n=66; epicardial, n=46) and 12 different studies downloaded from GEO [35] or ArrayExpress [36] (accession numbers: E-MTAB-1895, GSE20950, GSE26637, GSE27657, GSE27916, GSE27949, GSE40231, GSE41168, GSE66159, GSE71416, GSE82155, GSE9624; Supplementary Data S2).

Raw data (CEL-files) were downloaded and preprocessed together with the reference dataset as described above. Subsequently, CIBERSORT was used together with our custom AT21 signature matrix to deconvolute the 779 samples, determining their relative cell type composition. This allowed us to estimate the average cell type composition of four different adipose tissue depots (Figure 5) and to investigate associations between differences in adipose tissue cell type composition and various phenotypes (Figure 6).

Similarly, the deconvolution approach was applied to isolated cell types (reference dataset), to check how well closely related cell types (e.g. subcutaneous and pericardial adipocytes) can be distinguished (Figure 3C).

##### Statistics

Due to skewed distributions of relative cell type composition and a limited range of possible values between 0 and 1, we used non-parametric (Wilcoxon) tests to evaluate the significance of differences between phenotypes. Repeated measures within the same person (i.e. intervention studies with placebo, resveratrol, or calorie restriction) as well as the twin study (heavier vs. leaner twin concordant, heavier vs. leaner twin discordant) were considered as paired samples and analyzed using Wilcoxon signed rank test. Non-paired comparisons were performed using Wilcoxon rank sum test. Continuous analyses testing the association of age and BMI with adipose tissue cell type composition were performed using permutation tests based on Spearman correlation (‘cor.test’ function of package ‘stats’ in R). We report significant results both with and without Benjamini-Hochberg correction for multiple testing for the 336 comparisons (21 cell types times 16 phenotypic traits). Reported p-values are without correction.

#### Cell type-specific marker detector (CellMaDe)

A classical approach to cell type identification is the use of antibodies for specific marker proteins in immunohistochemistry or flow cytometry-based approaches. For these approaches, it is usually necessary to know cell type-specific markers that are not expressed (or only much lower expressed) in any of the other cell types. We refer to these markers as ‘primary markers’ in the manuscript. This approach comes with the limitation that some cell types are difficult to distinguish based on the expression of single marker proteins. For instance, mesenchymal stromal cells are typically characterized by a combination of several markers as well as functional assays [37]. Thus, where primary markers are not applicable, the idea is to combine several ‘secondary markers’ to receive unambiguous cell type identification.

In CellMaDe, we define the ‘**primary criterion**’ and the ‘**secondary criterion**’ to determine primary and secondary markers, respectively, as follows:

For each gene and each cell type, the primary criterion is calculated as the average expression of that gene in this cell type, minus the largest average expression of that gene in any other cell type, i.e.

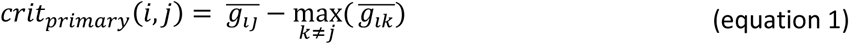

where 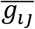 is the average expression of gene *i* in cell type *j*. The secondary criterion is calculated for each gene and each cell type as the average expression of that gene in this cell type minus the average expression of that gene in all other cell types, i.e.

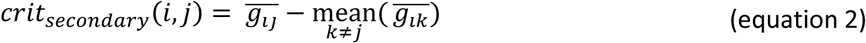

This results in a primary criterion score and a secondary criterion score for each gene which are then used to obtain the ranked lists for each score (Supplementary Figure S2).

##### Identification of cellular compartments

We have also annotated the top ten primary markers and selected conventional markers with their corresponding Gene Ontology (GO) cellular locations [38] in Fig 2b. Since these markers are identified to be used potentially in cell sorting, identification, or tracking applications, they are usually desired to be either on the cell surface, membrane or on the extracellular part of the membrane. Hence, we have used four different notations depicting the locations of interest (M: membrane, E: extracellular, ME: both membrane and extracellular, or O: other) for the corresponding proteins (Figure 2B, Supplemental Figure S2).

#### Evaluation of primary markers via Anatomically-annotated Tissue Expression Profiles

The definition of primary and secondary criteria defined in CellMaDe, depends on the cell types included in the analysis arguably and therefore, can be considered (adipose) tissue-specific, provided that all relevant cell types from the given tissue are included. In order to evaluate the validity of the most promising primary marker identified per cell type, as well as its general applicability across different tissues, we used Genevestigator [39] to compare the expression of this marker to a large compendium of different cell and tissue types (394 anatomically annotated tissue expression profiles). Genevestigator is a tool, that allows access to a normalized and curated database of publicly available transciptomics profiles and permits reproducible data analysis. It is freely available for analyzing the anatomical tissue expression for candidate genes.

The result of the most promising markers per cell type is shown in Supplementary Figure S3 where the expression of each gene is shown in a compendium of different tissue types. For the displayed figures, scatterplot and list options are selected with a log2 scale. All genes yielded results from all 394 anatomically annotated expression profiles except for the CellMaDe predicted primary marker Platelet Factor 4 Variant 1 (PF4V1) that is predicted for platelets. The expression of PF4V1 was only present in 48 anatomical tissues from the Genevestigator database where platelets were not present.

#### Independent (ex-vivo) validation of the TissueDecoder Framework

We performed an independent cross-platform validation of the TissueDecoder framework by integrating two datasets (GSE73174 and GSE80654, Affymetrix Human Transcriptome Array 2.0) containing samples from eight isolated cell types (CD4^+^ T cells, CD8^+^ T cells, CD14^+^ Monocytes, CD19^+^ B cells, CD56^+^ Natural Killer Cells – isolated from blood - and adipocytes, progenitors/adipose stem cells (CD45^-^CD34^+^CD31^-^), monocytes/macrophages (CD45^+^CD14^+^) – isolated from adipose tissue, Supplementary Data S2). The combined dataset is referred to as the “validation dataset”.

The two datasets were downloaded (CEL files) and preprocessed together as described in section deconvolution of adipose tissue cell types. Subsequently, we performed probe matching via the biomaRt R package for platform transformation and quantile normalized the validation dataset with the reference and analysis datasets form the Affymetrix Human U133 Plus 2.0 microarray.

The TissueDecoder framework is being used to calculate the percentages of the 21 cell types from the AT21 signature matrix in the validation dataset and to evaluate the expression of conventional markers as well as the primary markers reported from CellMaDe. The results are shown in Supplementary Figure S4 and Supplementary Data S2.

#### Literature review of adipose tissue cell type composition

We performed a literature review of quantitative reports about the cellular composition of the human adipose tissue for comparison with our results. Using a manual (non-systematic) search, we have extracted original and review articles, which we then further examined for relevant referenced studies. We selected a total of 22 original studies (Supplementary Data S4) that report cell type-specific cell counts in human adipose tissue (SAT and/or OAT), from which we extracted information about the cell type, number of people included with information on gender and BMI, adipose tissue depot, biopsy type, method and marker used for cell counting, as well as mean, standard deviation/standard error, minimum and maximum of reported counts (where available). In many studies, cell counts were only reported graphically, in which cases we extracted them manually through visual inspection.

Fifteen out of the twenty-two studies reported macrophage counts, of which eleven exclusively focused on macrophages. Frequencies of CD4^+^ T cells, CD8^+^ T cells, B cells, and endothelial cells are reported in three studies each. Natural killer cell frequencies are reported in two studies and frequencies of myeloid dendritic cells, (subcutaneous) adipocytes, and fibroblasts are mentioned only in one study each. Reports of adipose stem/stromal cell frequencies are included in six of the twenty-two studies. We have made the effort to compile the literature to the best of our knowledge, however, our list may not be exhaustive since several studies that focus on the isolation (rather than quantification) of adipose stromal cells also report cell numbers. We did not find any cell counts for the remaining eleven cell types in our signature matrix (monocytes, plasmacytoid dendritic cells, eosinophils, neutrophils, platelets, erythroblasts, smooth muscle cells, chondrocytes, osteoblasts, mesenchymal stromal cells, and pericardial adipocytes).

Direct comparison of the different studies is complicated not only by heterogeneity in the methods and markers used for cell counting but also by the heterogeneity in the units in which cell counts were expressed. The 22 studies used a total of six different units, namely (i) number per 100 adipocytes, (ii) number per g of adipose tissue, (iii) percent of SVF cells, (iv) number per total number of nuclei, (v) number per high power field, and (vi) number per mm^2^. We converted all units into percent of total cells using the formulas described below, excluding the two studies expressing the cell numbers in number per high power field and number per mm^2^.

##### Unit conversion

Two studies reported cell counts (macrophages) as number per total number of nuclei, counted via immunohistochemistry on tissue slices. We assumed that the number of anucleated cells in adipose tissue is negligible and directly used the reported number as the percent of total cells.

The unit number per 100 adipocytes was used in six of the included studies that determined macrophage frequency via immunohistochemistry. We converted the reported number (*x*) into percent of total cells (*y*) via the formula

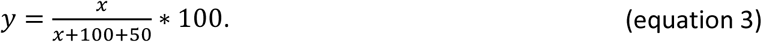

Here, *x*+100+50 represents the estimated number of total cells per 100 adipocytes, assuming that there are 50 cells other than adipocytes and macrophages, such as ASCs, other immune cells, and endothelial cells, per 100 adipocytes. Hence, we assume that adipose tissue roughly consists of 2/3 adipocytes and 1/3 other (SVF) cells, as long as the number of macrophages is not significantly high [40].

Four studies reported cell numbers in the unit number per g of adipose tissue based on flow cytometry or enzymatic isolation. We converted the reported number (*z*) into number per 100 adipocytes (*x*) via the formula

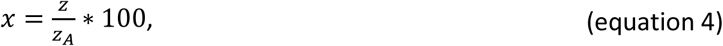

where *z_A_* is the number of adipocytes per g of adipose tissue. Subsequently, we used the formula described above to convert *x* into percent of total cells (*y*). We estimated *z_A_* as

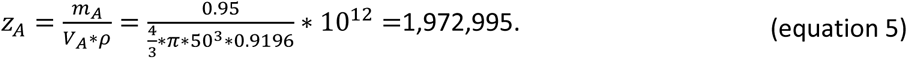

Here, *m_A_* = 0.95g is the assumed weight of adipocytes per gram of adipose tissue, *V_A_* is the average volume of adipocytes assuming that adipocytes are spherical with a radius of 50μm (diameter of 100μm) [41], and 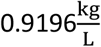 is the density of fat.

Finally, there were ten studies that used flow cytometry of the SVF to determine cell numbers as a percent of SVF. We converted this number to a percent of total cells through division by three, assuming that adipose tissue consists of roughly 1/3 SVF cells, as described above.

## List of abbreviations

SAT: Subcutaneous Adipose Tissue
OAT: Omental Adipose Tissue
EAT: Epicardial Adipose Tissue
PAT: Pericardial Adipose Tissue
SVF: Stromal Vascular Fraction
ASC: Adipose tissue Stem Cell
CAD: Coronary artery disease
CHD: Congenital heart disease

## Declarations

### Authors’ contributions

M.L. and G.E. conceived the project, performed the analyses and wrote the paper (all in equal contribution). All authors were involved in interpretation of results and editing of the paper.

